# Unraveling Tumor-Immune Heterogeneity in Advanced Ovarian Cancer Uncovers Immunogenic Effect of Chemotherapy

**DOI:** 10.1101/441428

**Authors:** Alejandro Jiménez-Sánchez, Paulina Cybulska, Katherine Lavigne, Tyler Walther, Ines Nikolovski, Yousef Mazaheri, Britta Weigelt, Dennis S. Chi, Kay J. Park, Travis Hollmann, Dominique-Laurent Couturier, Alberto Vargas, James D. Brenton, Evis Sala, Alexandra Snyder, Martin L. Miller

## Abstract

In metastatic cancer, the role of heterogeneity at the tumor-immune microenvironment, its molecular underpinnings and clinical relevance remain largely unexplored. To understand tumor-immune dynamics at baseline and upon chemotherapy treatment, we performed unbiased pathway and cell type-specific immunogenomics analysis of treatment-naive (38 samples from 8 patients) and paired chemotherapy treated (80 paired samples from 40 patients) high-grade serous ovarian cancer (HGSOC) samples. Whole transcriptome analysis and image-based quantification of T cells from treatment-naive tumors revealed ubiquitous variability in immune signaling and distinct immune microenvironments co-existing within the same individuals and within tumor deposits at diagnosis. To systematically explore cell type composition of the tumor microenvironment using bulk mRNA, we derived consensus immune and stromal cell gene signatures by intersecting state-of-the-art deconvolution methods, providing improved accuracy and sensitivity when compared to HGSOC immunostaining and leukocyte methylation data sets. Cell-type deconvolution and pathway analyses revealed that Myc and Wnt signaling associate with immune cell exclusion in untreated HGSOC. To evaluate the effect of chemotherapy on the intrinsic tumor-immune heterogeneity, we compared site-matched and site-unmatched tumors before and after neoadjuvant chemotherapy. Transcriptomic and T-cell receptor sequencing analyses showed that site-matched samples had increased cytotoxic immune activation and oligoclonal expansion of T cells after chemotherapy, which was not seen in site-unmatched samples where heterogeneity could not be accounted for. These results demonstrate that the tumor-immune interface in advanced HGSOC is intrinsically heterogeneous, and thus requires site-specific analysis to reliably unmask the impact of therapy on the tumor-immune microenvironment.

## INTRODUCTION

It is unclear how the complex interplay between tumor cells and the tumor microenvironment (TME) and their interactions affect treatment outcome in metastatic cancer (Kitamura, Qian, and Pollard 2015; Janssen et al. 2017; Robinson et al. 2017). Investigating this interplay in an advanced disease setting is complicated by the difficulty of obtaining multiple-site tumor samples and the finding that different tumors within the same individual can harbor distinct immune microenvironments (Sridharan et al. 2016; Jiménez-Sánchez et al. 2017a; Reuben et al. 2017; A. W. Zhang et al. 2018). Moreover, interactions between different cell populations of the TME are plastic and can change dependent on extrinsic perturbations such as therapy (Wang et al. 2016). In systems biology, perturbations are often used to infer how individual components of the system are interconnected by analyzing their dynamic behaviour in response to the external stimuli (Aldridge et al. 2006; Geva-Zatorsky et al. 2010; Molinelli et al. 2013). By analogy, immuno-oncology research needs to systematically characterize how oncogenic signaling activity affects the TME and immune infiltration, as well as to explore the variability and plasticity observed across clinical settings such as disease stage, anatomic sites and effect of treatment.

Heterogeneity in cancer spans multiple dimensions: at the molecular level with the presence of genetic intra-tumor heterogeneity (ITH) (Shah et al. 2009; Campbell et al. 2010; Gerlinger et al. 2012), the cellular level with variability observed in infiltration and recruitment of non-tumor cell populations in the TME (Natrajan et al. 2016), and the spatial and population levels where variability is observed both within tumors of the same individual with disseminated disease (Jiménez-Sánchez et al. 2017) and between tumors of different patients (McPherson et al. 2016; Patch et al. 2015). Thus, unbiased and systematic analysis of ITH and TME heterogeneity poses significant challenges. While the study of ITH has been facilitated by computational approaches to estimate the distribution and co-existence of different tumor clones from large-scale somatic mutation data, bioinformatics analysis of TME heterogeneity poses considerable technical difficulties due to the diversity of cell types present in the TME, in particular when distinguishing tumor from non-tumor cell populations based on bulk tumor analysis. Recent genomic approaches to characterize the TME include the development of immune cell deconvolution methods that aim to decompose a mixture of immune cell gene transcripts from bulk expression data (Finotello and Trajanoski 2018). However, no objective side-by-side comparisons of these methods have been implemented (Zheng 2017), and the accuracy and utility of immune cell deconvolution methods for determining TME heterogeneity in clinical settings is uncertain.

High grade serous ovarian cancer (HGSOC) is ideally suited to the study of TME heterogeneity owing to its clinical presentation with multisite abdominal disease and standardized treatment with either optimal surgical debulking at diagnosis or delayed primary surgery after neoadjuvant chemotherapy (NACT) (Bowtell et al. 2015). Thus, in HGSOC there is a unique opportunity to study the characteristics of the TME at multiple sites and to observe variation at baseline (diagnosis) and following perturbation with platinum-based chemotherapy, with the underlying hypothesis that the role of the TME may be exposed by its dynamic response to extrinsic perturbations (e.g. chemotherapy). Furthermore, since HGSOC is typically diagnosed when dissemination has already taken place in the peritoneal cavity, this malignancy provides the basis for evaluating the ubiquitousness of intra-patient TME heterogeneity in an advanced, metastatic disease setting. In HGSOC, the low somatic point mutation load, high aneuploidy levels and high copy number alterations have been associated with lack of immunogenicity (Bowtell et al. 2015; A. W.Zhang et al. 2018). Despite the low intrinsic immunogenicity, T cell infiltration plays a major role in predicting HGSOC survival in a primary disease setting (L.Zhang et al. 2003; Ovarian Tumor Tissue Analysis (OTTA) Consortium et al. 2017), and recent studies have started to shed light on the interplay between ITH and T cell interactions (A. W.Zhang et al. 2018), as well as the potential effect of chemotherapy on T cell infiltration in HGSOC (Böhm et al. 2016). However, the extent of TME heterogeneity has not been systematically characterized in metastatic disease, including advanced HGSOC, and its underlying mechanisms and role in therapeutic response remain unknown.

To characterize heterogeneity at the level of the TME and to begin to identify the molecular and cellular underpinnings of immune infiltration variability at diagnosis and after perturbation by chemotherapy, we performed a systematic analysis of >100 HGSOC samples from treatment-naive and NACT patient cohorts. Our findings confirm that HGSOC is a disease characterized by pervasive TME heterogeneity with distinct immune microenvironments co-existing in different tumor nests within the same individuals at diagnosis. We leverage our rich data sets to create an ensemble computational approach that integrates and improves upon existing immune and stromal cell deconvolution methods, thus enabling us to systematically characterize the TME of HSGOC before and after treatment. We identify oncogenic signaling pathways such as Myc and Wnt that associate with immune cell exclusion when comparing tumors with high cancer cell fraction (high purity) vs low cancer cell fraction (low purity). We find that NACT induces immune activation and specific T cell clonal expansions in local TMEs, however, intra-patient TME immune heterogeneity can mask such effects. Consequently, systemic immunomodulatory therapies may be ineffective in a subset of tumor sites, thus preventing overall patient benefit. Together, these results show that intra-patient TME heterogeneity is ubiquituos in HGSOC, which could confound clinical outcomes.

## RESULTS

### Intrapatient Transcriptomic Heterogeneity is Largely Explained by Immune Signaling

To investigate the tumor microenvironment (TME) of HGSOC in a treatment-naive context, we analyzed the transcriptome of 38 primary and metastatic tumor samples from 8 treatment-naive patients collected prospectively (Figure 1A, Supplementary Table 1A and B). Primary tumor masses and peritoneal metastases were resected and placed on patient-specific 3D moulds created based on tumor segmentation using high resolution T2-weighted magnetic resonance (MR) images [REF:Weigelt B, Vargas AH, Selenica P, Geyer FC, Mazaheri Y, Blecua P, Conlon N, Hoang LN, Jungbluth AA, Snyder A, Ng CKY, Papanastasiou AD, Sosa RE, Soslow RA, Chi DS, Gardner GJ, Shen R, Reis-Filho JS, Sala E. Radiogenomics analysis of intra-tumor heterogeneity in high-grade serous ovarian cancer. BJC (under review)]. Each specimen was placed in the custom-made 3D mould in the operating theatre and was further dissected into sub-specimens according to three multi-parametric imaging-based phenotypically distinct clusters, hereafter referred to as “habitats”. Habitats were obtained from MR and 18F-FDG-PET imaging and were defined based on quantitative imaging features that measure water diffusion, micro-capillary perfusion, permeability and metabolic activity (see Methods). We first performed an unbiased clustering analysis of the whole transcriptome. We observed that overall gene expression of tumor samples was highly patient specific, irrespective of anatomical site using t-distributed stochastic neighbor embedding (t-SNE) which accounts for nonlinear relationships (Figure 1B). To focus on well-defined biological processes and signaling pathways, we performed ssGSEA (Hänzelmann, Castelo, and Guinney 2013) using the hallmark gene sets (Arthur Liberzon et al. 2015), stromal and immune gene signatures, and tumor cell fraction (purity) using the ESTIMATE algorithm (Yoshihara et al. 2013). We categorized the gene sets into five classes: oncogenic, cellular stress, immune, stromal, and other. Principal component analysis (PCA) showed that most of the gene set expression variation between samples (60% of variation) could be explained by oncogenic, immune, and stroma-associated gene sets (Figure 1C, S1A). In contrast to the full transcriptome analysis, the patient specific clustering was less evident, indicating that tumors from different patients share common patterns of pathway activation and non-cancer cell infiltrates. To investigate which gene sets explained most of the observed variance, we computed the principal component feature loadings and displayed them in a variable factor map (Figure 1D). This analysis showed that PC1 (40% of variation) is largely explained by tumor purity, since immune and stromal vectors had an opposite direction to oncogenic vectors and tumor purity (immune vs oncogenic: *p*=3e^-05^; stromal vs oncogenic: *p*=1.3e^-03^), and PC2 (20% of variation) further showed a separation of immune, stromal, and cellular stress vectors (immune vs stromal: *p*=0.046; immune vs stress: *p*=0.041; Figure S1B).

**Figure 1:**
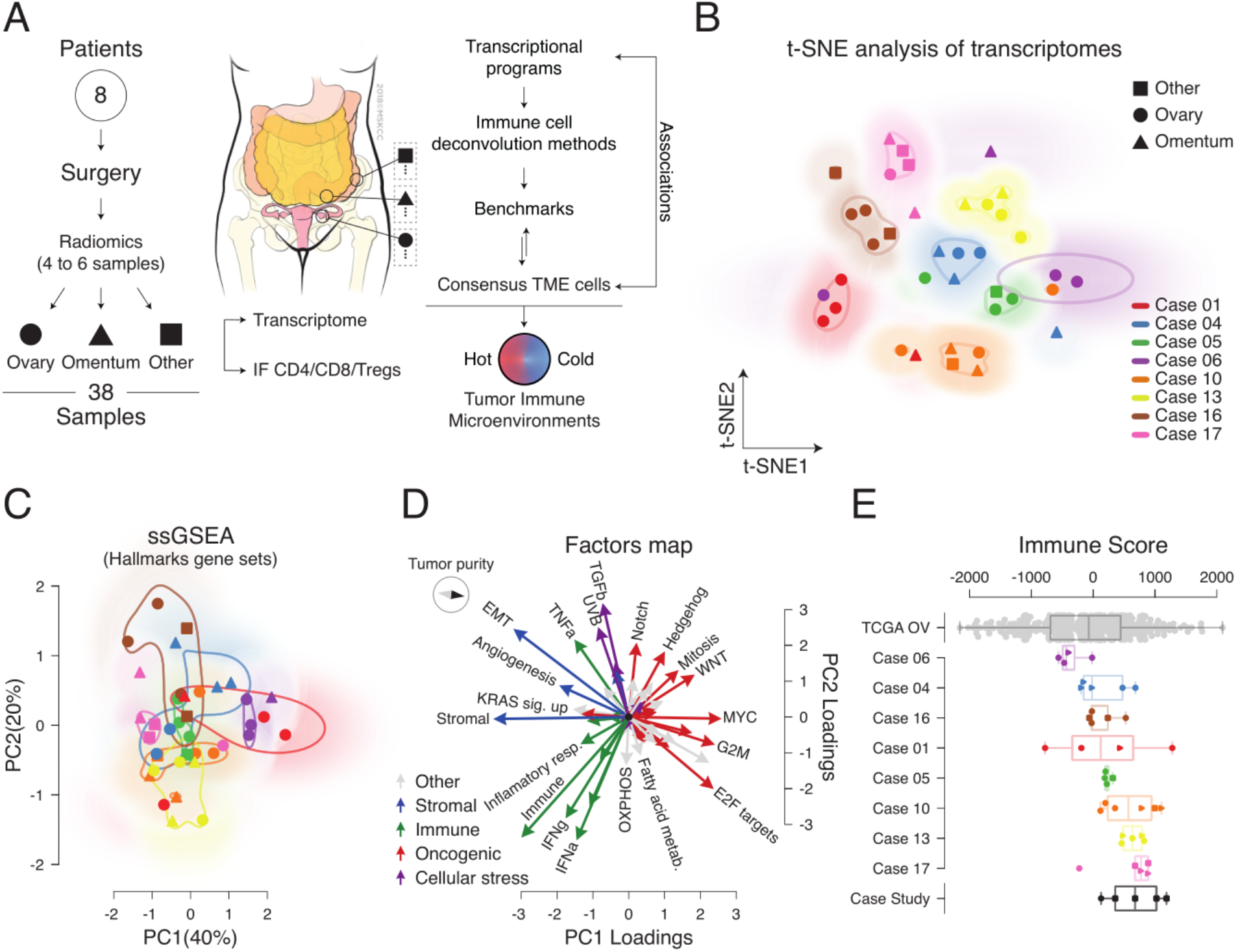
Immune signaling contributes to the majority of the transcriptional variance observed across multiple tumor samples from treatment-naive HGSOC patients. A) Flowchart of sample acquisition and analysis. Peritoneal metastases other than omentum were defined as “Other”. B) t-SNE analysis of overall transcription profiles of multiple HGSOC tumor samples per patient. C) PCA of ssGSEA-based analysis of hallmark gene sets. D) Principal component feature loadings (magnitude and direction) of C are shown in the variables factor map. Vectors are colored according to a major biological classification of hallmark gene sets. Variation across classes in PC1 (*p*=3.2e^-16^) and PC2 (*p*=0.02) after Kruskal-Wallis H-test (Figure S1). Directionality of ESTIMATE’s tumor purity is represented with the map compass. E) ESTIMATE immune score across patients and samples. The Case Study samples were taken from (Jiménez-Sánchez et al. 2017). The bottom and top edges of the box plots indicate the 25th and 75th percentiles.

Since immune related pathways explained a significant amount of variation between the samples, we further investigated the extent of intra-patient immune heterogeneity by computing the ESTIMATE immune score for each sample. In addition, we included as a reference the immune scores of the samples from a HGSOC case study with >9 years of clinical history we previously analyzed (Jiménez-Sánchez et al. 2017) and the immune scores of ovarian cancer samples from The Cancer Genome Atlas (TCGA), which comprises 307 treatment-naive primary tumors (Cancer Genome Atlas Research Network 2011). Overall, the immune scores of our cohort fell within the range expected at the population level (Figure 1E). Some patients (01, 04, 10, and the case study) showed an intra-patient variation comparable to the inter-patient variation observed at the population level by the TCGA ovarian cancer samples, which indicates that within a single individual, complete distinct immune microenvironments can co-exist at diagnosis of HGSOC. Also, all patients in the cohort had at least one sample with similar or lower immune score than the progressing and immune excluded tumors of the case study, where distinct tumor-immune microenvironments led to different clinical outcomes (Jiménez-Sánchez et al. 2017). Importantly, consistent with our prior report, we recapitulate the observation that tumors with high immune signaling and immunosuppressive Wnt signaling tend to be mutually exclusive (Figure 1D).

### Co-existence of distinct tumor-immune microenvironments in treatment-naive HGSOC

To further characterize the tumor microenvironment of HGSOC, we performed multicolor immunofluorescence (IF) staining and quantification of CD4, CD8, and regulatory T cells (CD4+ FOXP3+) in at least 10 tumor regions excluding stromal areas in each sample leading to a compendium of 440 imaged and digitally quantified tumor regions (Figures 2A-B, S2, and Supplementary Table 2A). This multi-region and multi-site IF analysis shows that treatment-naive HGSOC patients present variation in T cell infiltration in tumor deposits, ranging from less than 1% (e.g. patient 6) to T cells accounting for more than 10% of total cells in some areas (e.g. patient 01 and 10). Furthermore, some patients’ tumor deposits demonstrated marked variation in T cell infiltration within the same tumor deposit across different habitats (e.g. patient 01). We then performed a linear mixed effects model analysis (see Methods and Supplementary Table 2B) to statistically evaluate whether there is a difference of T cell infiltration between patients, between tumors of the same patients and between habitats within tumors. We found a remarkable difference in T cell infiltration across tumors within patients (Degrees of freedom=2, CD8 F-value=8758; CD4 F-value=58; Tregs F-value=657) and habitats within a tumor (Degrees of freedom=2, CD8 F-value=1184; CD4 F-value=2870; Tregs F-value=2216). Sites and habitats also showed an important level of interaction (Degrees of freedom=4, CD8 F-value=5915; CD4 F-value=4466; Tregs F-value=142). This systematic T cell IF staining and computerized cell detection-counting confirm the variation in T cell infiltration across patients, across tumor samples within patients, within tumors and within habitats. Together with the transcriptome analysis, these data show that HGSOC is intrinsically characterized by the presence of heterogeneous immune TMEs within patients and within tumors.

**Figure 2.**
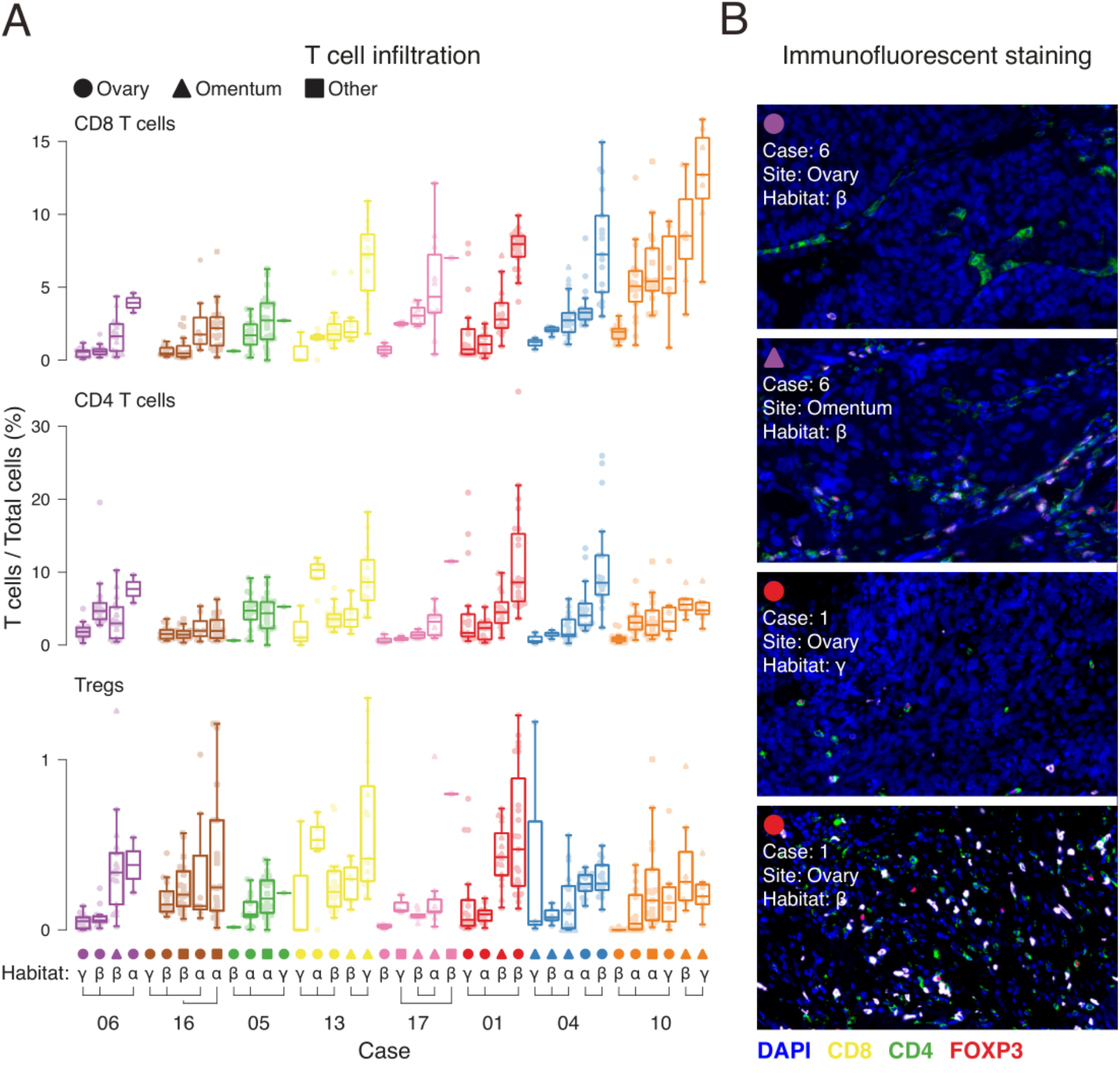
T cell infiltrate variation across patients, within patients, and within tumors. A) Multi-tumor sampling from 8 HGSOC patients are shown with each dot representing the percentage of T cell subsets in a quantified area within a given tumor section stained with multicolor IF for CD8, CD4 and FOXP3. Stromal areas were excluded based on H&E stains. Patient cases are indicated by different colors. Anatomical sites of tumor deposits are indicated by different markers (circle, triangle and square). Habitats are defined by the greek letters α, β and γ. Habitats from the same tumor are indicated by connecting lines. Boxplots are sorted according to the median of CD8 T cell infiltration across patients, sites and habitats accordingly. B) Representative images of panel A.

### Consensus tumor microenvironment gene sets improve cell deconvolution from bulk tumor mRNA

To estimate the relative abundance of different cell types in an unbiased manner using bulk RNA data, various computational approaches have been developed during the last decade (Finotello and Trajanoski 2018). However, a comparison that objectively evaluates the performance of these approaches against one another has not been conducted and no publicly available data sets have been generated to serve as ground truths thus far (Zheng 2017). Therefore, to test the performance of deconvolution methods, we performed a preliminary benchmark using the T cell IF quantification of the 440 regions from the 38 treatment-naive HGSOC samples as the ground truth. We used ESTIMATE for total T cell infiltration (Yoshihara et al. 2013), and compared CIBERSORT (Newman et al. 2015), TIMER (B. Li et al. 2016), MCP-counter (Becht et al. 2016) and xCell (Aran, Hu, and Butte 2017) for cell type specific deconvolution. We also evaluated immune gene sets that were defined based on gene expression of sorted immune populations (Bindea et al. 2013), as well as immune gene sets based on the Immunological Genome Project database (Davoli et al. 2017; Heng, Painter, and Immunological Genome Project Consortium 2008). Using ssGSEA (Hänzelmann, Castelo, and Guinney 2013), the Bindea et al. and Davoli et al. gene sets were used to calculate normalized enrichment scores (NES) of the corresponding cell types. To be able to evaluate the performance of ESTIMATE, which calculates a total immune score but does not deconvolute immune cell types, total immune scores from the deconvolution tools were generated (see Methods). Total immune scores and deconvolution scores for CD4, CD8 and Tregs were correlated independently against the aggregated T cells and the corresponding cell type fractions (Figure 3A). Importantly, not all of the methods selected deconvolute CD4, CD8, and Tregs; however, for the methods that deconvolute all three cell types, none consistently outperformed the other methods in this independent benchmark analysis. In addition, none of these methods was able to get a significant positive correlation with Tregs, which may indicate a lower-limit threshold of sensitivity of detection, since Tregs comprised, on average, less than 1% of cells in the tumor nests (Figure 2A, S2).

**Figure 3.**
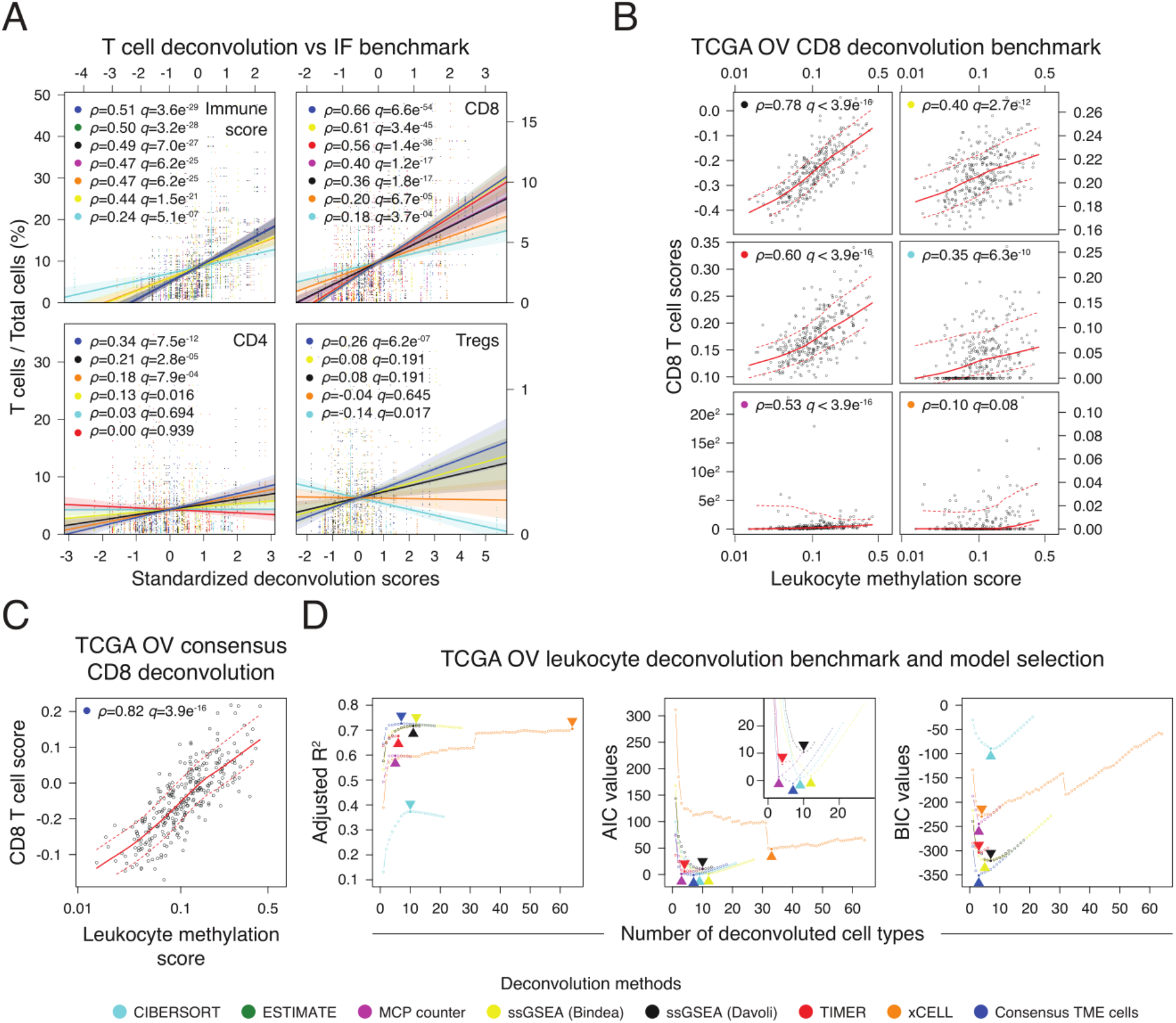
Consensus tumor microenvironment cell deconvolution method improves estimation of tumor infiltrating T cells and leukocytes. A) Spearman’s rank-order correlations between percentage of T cells (immunofluorescent staining) and the corresponding deconvolution scores for each method using the treatment-naive ovarian cancer samples. Correlation coefficients (*ρ*) and *q*-values are ordered from higher and more significant to lower and less significant. Deconvolution scores were standardized (z-score) to visually compare the different correlations in the same scale. IF immune score was calculated by adding up CD8, CD4, and Treg counts as an approximation, while deconvolution immune scores were calculated according to each deconvolution tool (see Methods). B, C) TCGA ovarian cancer Spearman’s rank-order correlations between CD8 T cell deconvolution scores and leukocyte methylation scores. D) Multiple linear regression analysis using leukocyte methylation score as response variable, and deconvoluted cell types as explanatory variables (see Methods). Adjusted R2, Akaike information criterion (AIC), and Bayesian information criterion (BIC) values were calculated to compare goodness of fit and model simplicity. Arrowheads indicate best model for each method. Inset in the AIC panel shows a magnification of the best ranked models.

Since the cell type deconvolution methods were developed independently of each other, we reasoned that generating Consensus gene sets by including the genes that fall in the intersection of different cell types across the different tools could improve the cell deconvolution performance. Since not all methods deconvolute the same cell types, we focused on cell types that at least two methods deconvolute. We selected genes that overlapped between the independent methods (intersection), and finally removed genes whose expression levels positively correlated with tumor purity using TCGA ovarian cancer samples as a reference (see Methods and Figure S3A). We correlated the ssGSEA NES of the Consensus TME cell gene sets against the fraction of T cells quantified, and observed that the Consensus gene sets consistently showed higher positive correlations than the individual methods. In addition, the consensus Tregs NES was the only score with a significant positive correlation with the fraction of Tregs, suggesting a greater level of sensitivity obtained with the Consensus approach (Figure 3A, Spearman’s rho=0.26, *q*=6.2e^-07^).

To further benchmark the methods and the consensus signatures, we employed TCGA ovarian cancer leukocyte methylation scores (Cancer Genome Atlas Research Network 2011), which is an independent and larger patient cohort. Leukocyte methylation measurements provide orthogonal means toward estimating immune infiltration in tumors, and have been shown to significantly correlate with histological purity estimates in primary HGSOC (Carter et al. 2012). Importantly, the leukocyte methylation signature was generated by comparing methylation patterns between HGSOC tumors, normal fallopian tube samples and buffy coat samples of female individuals, making this data set ideal for benchmarking the deconvolution methods (Carter et al. 2012). We first performed a benchmark of all methods using the leukocyte methylation scores and the CD8 T cell deconvolution scores (Figure 3B), as i) this subpopulation is a major component of infiltrating leukocytes, ii) all deconvolution methods tested deconvolute CD8 T cells, and iii) overall the CD8 IF estimations were the best correlations (Figure 3A). Among the seven methods, the consensus CD8 gene signature correlated best with the CD8 T cell score (Figure 3C, Spearman’s rho=0.82 *p*=3.9e^-16^). However, as the leukocyte methylation score does not count CD8 T cells exclusively, we then compared the different methods in an unbiased manner by fitting a multiple linear regression model for each method using the leukocyte methylation score as a response variable and the different cellular scores as explanatory variables, followed by unsupervised nested variable selection (see Methods and Figure S3B). We compared the proportion of leukocyte methylation score variance that is explained by the unsupervised selected deconvoluted cells (adjusted R-squared), as well as the relative quality of the models by considering goodness of fit and model simplicity (see Methods). The Consensus gene signatures provided the highest adjusted R-squared with fewer cell types selected (Adj. R-squared=0.73, *p*<2.2e-16, Figure 3D left panel), as well as being selected as the simplest and most accurate model to explain leukocyte methylation (Figure 3D middle and right panels). Finally, the consensus gene signatures that the systematic unbiased analysis suggested best explained leukocyte methylation were cells expected to be present in leukocyte infiltrates, with CD8 and NK cells accounting for the vast majority of the variation explained (Supplementary Table 3, CD8 *p*=1.74e^-07^, and NK cells *p*=1.79e^-11^). In addition, we performed a sensitivity analysis of the leukocyte methylation benchmark, and the consensus was also the best method with the same cells explaining leukocyte methylation selected (Figure S3C and Supplementary Table 3). Together, these benchmarks show a consistent improvement of leukocyte cell deconvolution provided by the Consensus gene sets in ovarian cancer samples.

### Tumor microenvironment cell deconvolution and molecular comparison of high and low purity treatment-naive HGSOC

Having generated our own robust method for immune cell deconvolution, the Consensus method, we applied the Consensus gene sets to the treatment-naive HGSOC transcript data to systematically assess if specific transcriptional programs were associated with variability in immune infiltration. We first visualized the variation across samples using the NES of deconvoluted Consensus gene sets of cells (Figure 4A, S4A). The gene sets explaining most of the variation were cytotoxic, NK cells and fibroblast being negatively correlated with tumor purity; while endothelial, monocytes and B cells positively correlated with tumor purity (Figure 4B, S4B). The NES of deconvoluted cells of this cohort were comparable to the NES obtained in the TCGA ovarian cancer data set (Figure S5). Interestingly, the cells with highest NES in most samples were fibroblasts, highlighting the intrinsic low-immunogenic nature of HGSOC (Wang et al. 2016).

**Figure 4:**
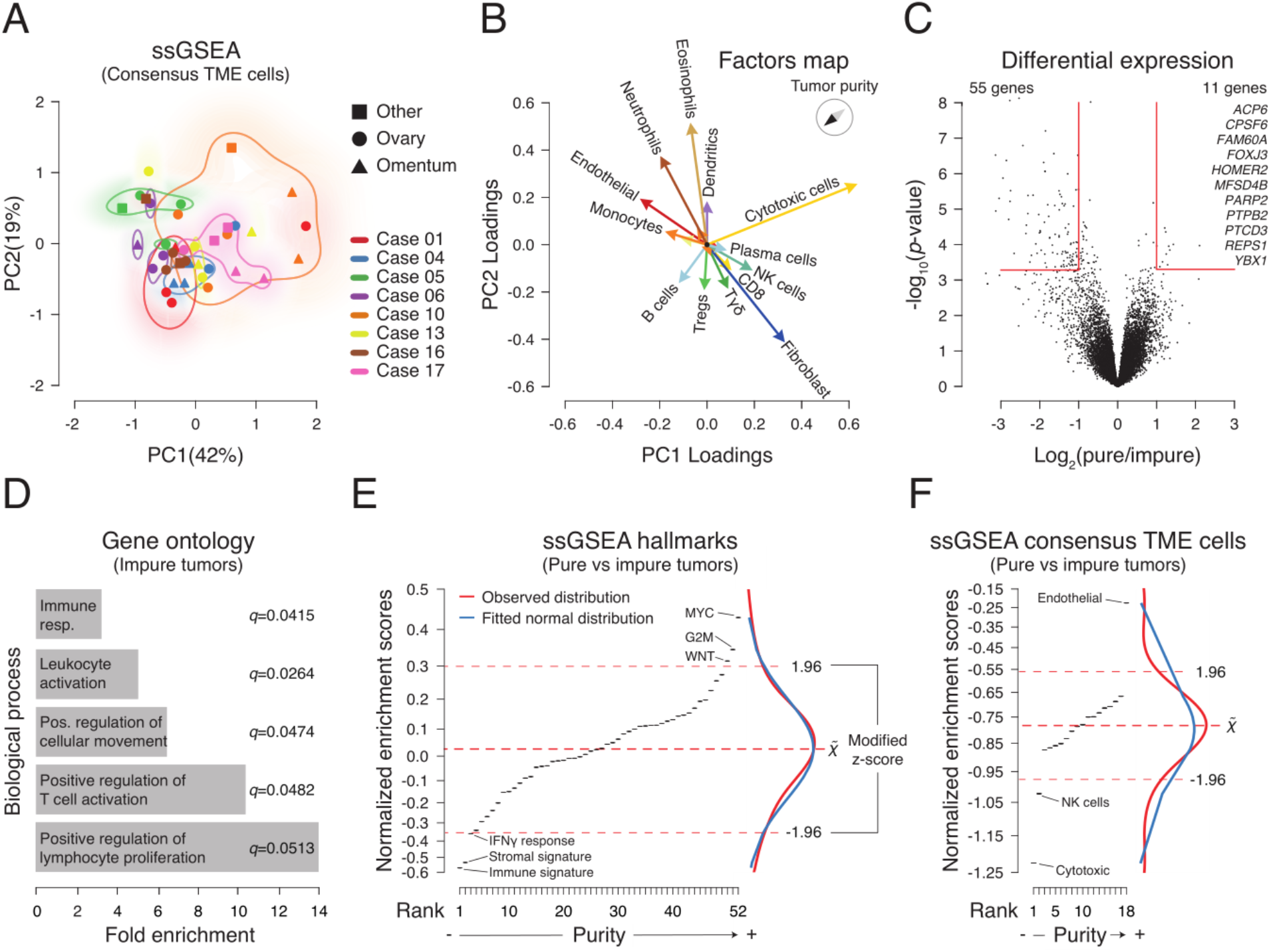
Unbiased analysis of tumor microenvironment heterogeneity in treatment-naive HGSOC tumors. A) PCA of ssGSEA-based analysis using the consensus deconvoluted cell gene sets. B) Principal component feature loadings (magnitude and direction) of A. Vectors are colored according to cell types, for example monocytes and macrophages M0, M1, M2 (orange), B cells and plasma cells (light blue), and CD8 and cytotoxic cells (yellow). C) Differential expression analysis of high purity and low purity classified tumors using the median purity score of the cohort as a cutoff (see Methods). Vertical red lines indicate +/- 1 fold change of gene expression, and the horizontal line indicates the corresponding 0.05 *q*-value on the y-axis. D) Gene ontology analysis of significantly highly expressed genes on low purity tumors. Significantly highly expressed genes in pure tumors are not significantly over represented in any gene ontology biological process. E, F) ssGSEA analysis of differentially expressed genes using hallmarks and consensus deconvoluted cell gene sets, respectively (see Methods). Gene sets on the x-axes were ranked according to their normalized enrichment score (Supplementary Tables 4A-B). Higher normalized enrichment score are indicative of higher purity scores. Dashed red lines indicate median and + / - 1.96 median absolute deviations (modified z-score) to define outliers. Marginal density plots of observed and values fitted to a normal distribution are shown.

To investigate genes associated with tumors with high cellularity (pure tumors), we used the median tumor purity of the cohort to classify high and low purity tumors (see Methods), and performed a differential expression analysis leveraging sample-patient dependency (i.e. considering that multiple tumors come from the same individual) to increase statistical power. As expected, genes related to immune activation were significantly highly expressed in low purity tumors, but only eleven genes were significantly highly expressed in the purer tumors compared to the lowly pure ones (Figure 4C). Gene ontology analysis (GO) showed that genes with significant higher expression in low purity tumors are significantly enriched in leukocyte proliferation and activation GO biological processes (Figure 4D), whereas no significant GO enrichment was found with the genes significantly highly expressed in pure tumors. The genes with significantly higher expression in pure tumors have been implicated in transcription [*CPSF6* (Rüegsegger, Blank, and Keller 1998), *FOXJ3* (Landgren and Carlsson 2004)], cellular growth [*HOMER2* (Tu et al. 1998; Xiao et al. 1998), *REPS1* (Cantor, Urano, and Feig 1995; Hu and Mivechi 2003)], glucose transport [*MFSD4B (Horiba et al. 2003)*], mitocondrial generation [*PTCD3 (Davies et al. 2009)*], aberrant proliferation [*YBX1* (Frye et al. 2009; Weidensdorfer et al. 2009)], ovarian cancer initiation and progression [*ACP6 (Hiroyama and Takenawa 1999; Fang et al. 2002)*, *PARP2 (Amé et al. 1999; Gunderson and Moore 2015)*] and TGF-beta signaling downregulation [*FAM60A (Muñoz et al. 2012; Smith et al. 2012)*], which consequently can directly and indirectly promote tumorigenesis through TME immunosuppression (Colak and ten Dijke 2017; Tauriello et al. 2018; Mariathasan et al. 2018).

To further investigate which molecular signaling pathways are more highly enriched in pure tumors, we performed ssGSEA using the adjusted *p*-values and changed the sign, positive or negative, according to the differential expression direction (see Methods). As expected, immune and stromal signatures were highly enriched in low purity tumors, in addition to IFN-gamma response. In contrast, Myc and Wnt signaling appeared to be highly enriched in pure tumors, both of which have been previously associated with immune exclusion in pre-clinical models of lung cancer (Kortlever et al. 2017) and melanoma (Spranger, Bao, and Gajewski 2015; Spranger et al. 2016; Spranger, Bao, and Gajewski 2014), respectively (Figure 4E). Not surprisingly, the proliferation related hallmark G2M was highly enriched in pure tumors. Of note, little or no overlapping between the G2M, Myc and Wnt hallmark gene sets was observed (Figure S4C and Supplementary Table 4A; 6 out of 258 genes overlapped between G2M and Myc, and 2 out of 242 genes overlapped between G2M and Wnt signaling gene sets). Considering the TME, cytotoxic and NK deconvoluted cells were preferentially enriched in low purity tumors, whereas endothelial cells were highly prevalent in pure tumors (Figure 4F). Since all genes in the cytotoxic gene set are included in the CD8 and NK cell gene set, this suggests that a particular activation of NK cells is more prominent in low purity tumors (Figure S4C), whereas pure tumors only showed enrichment of endothelial cells (Supplementary Table 4B). These observations suggest that Myc and Wnt signaling gene set enrichments in pure tumors could be considered at least partially independent of tumor proliferation, and may also contribute to immune cell exclusion as suggested by other studies (Spranger and Gajewski 2018).

### Chemotherapy induces immune activation in HGSOC

To investigate the effect of chemotherapy on the TME and evaluate if there is a confounding effect of the intra-patient TME heterogeneity described above, we studied the transcriptome of site-matched (n=18) and site-unmatched (n=38) primary and disseminated tumors before and after treatment with neoadjuvant platinum and taxane chemotherapy in 28 HGSOC patients (Figure 5A, Supplementary Table 5A-B). Using t-SNE dimensionality reduction on the whole transcriptomes, we found that treated and untreated samples cluster separately (Figure 5B), in contrast to the treatment-naive samples that cluster in a patient-specific manner (Figure 1B). Using the ssGSEA NES of the hallmark gene sets of site-matched and site-unmatched samples, we observed that treated and untreated sample groups were separated by the two first principal components with 63% and 50% of variation in site-matched and site-unmatched groups, respectively (Figures 5C and S6A-B). Both site-matched and site-unmatched groups showed that oncogenic and immune/stromal hallmarks contributed significantly to the variation explained by the first principal components (Figure S6C-D). However, only site-matched PC1 reached statistical significance after paired comparison between pre-and post-NACT samples (Figure S6E-F), while also explaining more than 50% of the variation in the site-matched samples (Figure S6A-B). Interestingly, cellular stress related pathways were more enriched in site-unmatched post-NACT than site-matched samples, potentially reflecting the cellular stress generated by therapy, whereas in site-matched samples, immune related pathways dominate the variation signal (Figures 5C, S6C-D). In addition, Wnt and Myc signaling showed a clear negative association to immune related gene sets in the site-matched samples, whereas no clear clustering contribution of Wnt and Myc gene sets is discernible in the site-unmatched samples (Figure 5C).

**Figure 5:**
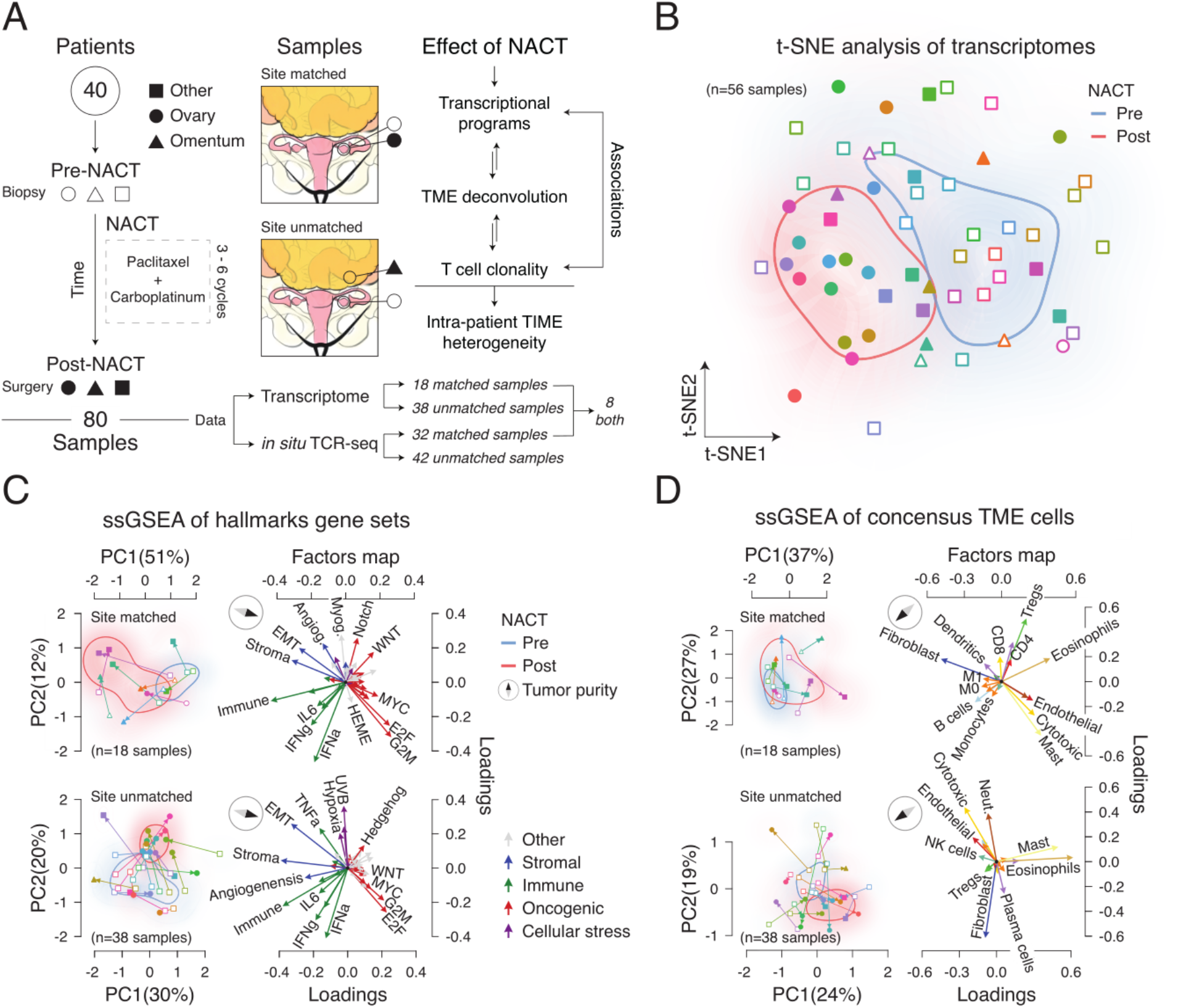
Unbiased signaling pathway and tumor microenvironment cell decomposition analysis of chemotherapy treated HGSOC site-matched and unmatched tumor samples. A) Flowchart of sample acquisition, clinical study design, and analysis. TME: Tumor microenvironment. B) t-SNE analysis of overall transcription profiles of multiple HGSOC tumor samples per patient. C, D) PCA and principal component feature projections (magnitude and direction) of ssGSEA-based analysis of hallmark gene sets and consensus tumor microenvironment (TME) cells respectively. Arrows in the principal component space indicate pre-to post-NACT directionality. Hallmark gene set vectors are colored according to a major biological classification. Angiog: Angiogenesis, Myog: Myogenesis. Consensus TME vectors are colored according to cell types. Neut: Neutrophils.

We then deconvoluted the TME cellular mixtures to investigate which cells were differentially present in the pre-and post-treatment tumors. Pre-and post-treatment samples also clustered separately both in site-matched and site-unmatched samples, with the first principal components accounting for 64% and 43% of variation explained, respectively (Figures 5D, S7A-B). Eosinophils and cytotoxic cells showed a negative association with tumor purity in site-matched samples, in contrast to fibroblasts, B cells and macrophages (Figures 5D, S7C). Interestingly, the only principal components that were significantly different between pre-and post-NACT samples were the first two principal components of the site-matched tumors (Figure S7E-F).

### Tumor-immune microenvironment intra-patient heterogeneity masks chemotherapy induced immune activation effect

To directly evaluate differences between pre-and post-treatment samples, we performed an exploratory data analysis leveraging the possibility of performing paired comparisons using the hallmark and consensus TME gene set NES independently for site-matched and site-unmatched samples (Figure 6A). Site-matched samples showed a clear increase of immune pathways and consensus TME gene sets in post-treatment samples, while site-unmatched samples showed an increase of cellular stress pathways reflecting cellular and metabolic stress after cytotoxic drug exposure, but no difference of consensus TME gene sets was detected in the site-unmatched cohort. Since we observed that cytotoxic and NK cell Consensus gene sets were mainly enriched in treatment-naive low purity tumors (Figure 4F), and it is known that CD8 T cells play a crucial role in ovarian cancer recurrence and overall survival (L.Zhang et al. 2003), we performed a multivariate Student’s t-statistic hypothesis test using cytotoxic, NK and CD8 T cell Consensus gene sets. We compared the difference between these 3 Consensus gene sets in pre-and post-NACT site-matched and site-unmatched samples (Figure 6B), and observed a significant increase of these immune cell types upon chemotherapy in the site-matched samples (*p*=0.0034), but no difference in the site-unmatched samples (*p*=0.92). To further evaluate T cell infiltration and activation between pre-and post-NACT samples, we performed *in situ* TCR sequencing. Since T cell activation leads to clonal expansion of particular T cell clonotypes, TCR clonality measures can be used as a surrogate for T cell activation upon specific (neo)antigen recognition (Pielou 1966; Kirsch, Vignali, and Robins 2015; Jiménez-Sánchez et al. 2017a). TCR clonal expansion was significantly higher in post-NACT site-matched samples (Figure 6C, *p*=0.001), but no significant difference was observed in site-unmatched samples (*p*=0.2). T cell fraction was also significantly higher in post-NACT site-matched samples (*p*=0.03), while a slightly lower T cell fraction was observed in site-unmatched post-NACT tumors, potentially as a result of the variability of immune infiltration between omentum metastases (pre-NACT biopsies) and primary tumors (post-NACT debulking surgery). To test whether the biopsy intervention could be a confounder and lead to an immune activation *per se*, we compared post-NACT site-matched and post-NACT site-unmatched TCR clonal expansions, since the post-NACT site-unmatched tumors were not originally biopsied. No significant difference was observed in TCR clonal expansion (*p*=0.67), suggesting that T cell clonal expansion was independent of biopsy treatment and likely induced by NACT. However, a significant increase of T cell density was observed in site-matched compared to site-unmatched post-NACT tumors (*p*=3.59e^-05^), potentially suggesting that wound healing after the biopsy procedure could increase the influx of T cells. Together, these results provide evidence that neoadjuvant chemotherapy induces an immune activation in the local TME of HGSOC, and that intra-patient inter-site TME heterogeneity can obscure this clinically relevant observation among tumor deposits within patients.

**Figure 6:**
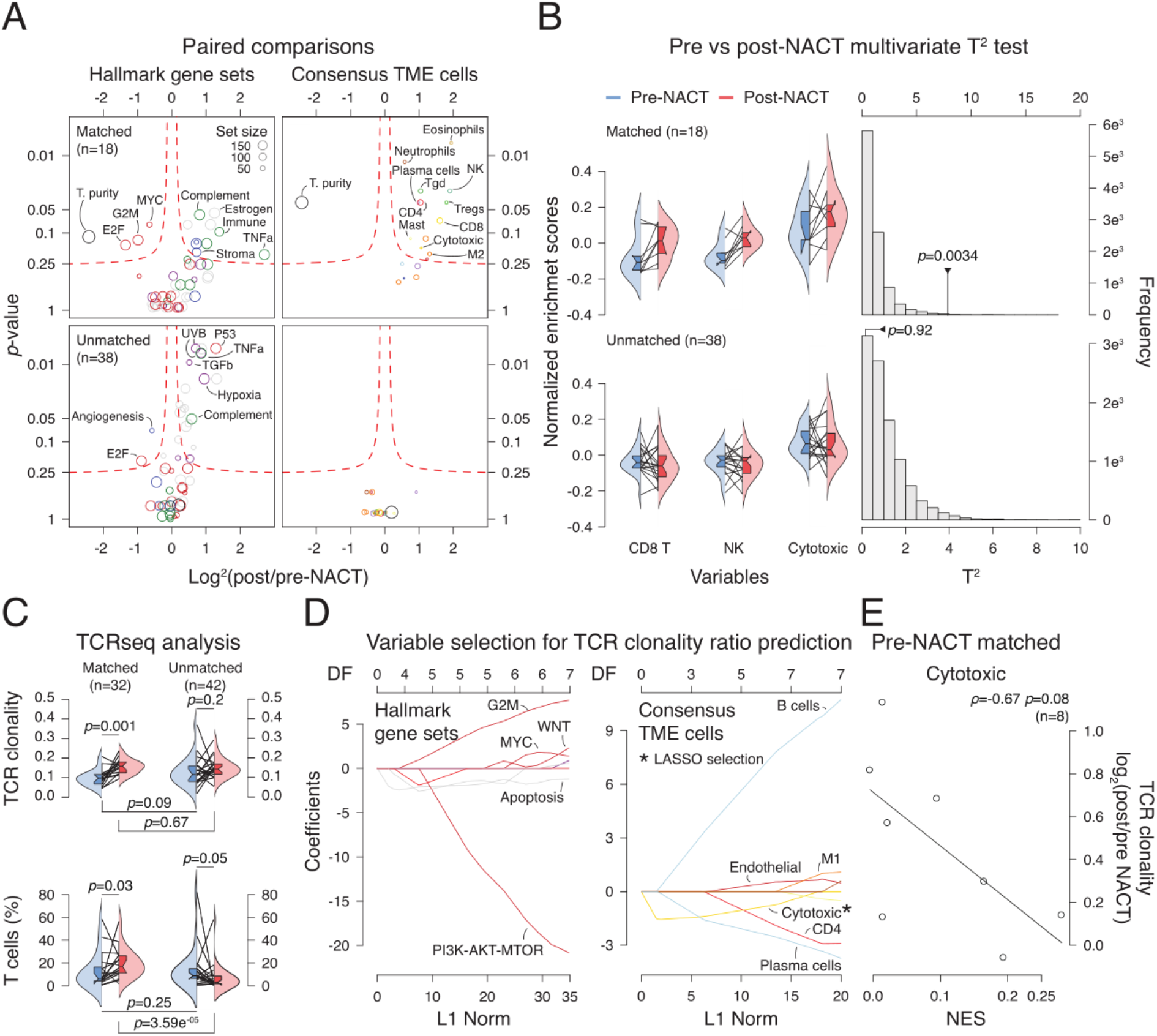
Immune activation induced by neoadjuvant chemotherapy is evident in site-matched but not site-unmatched sample analysis. A) Exploratory pre/post NACT paired comparisons of hallmark gene sets and consensus TME deconvoluted cells (see Methods). B) Multivariate T^2^ tests comparing pre and post-NACT CD8 T cells, NK cells, and cytotoxic NES together. C) Comparisons of TCR productive clonality (top), and percentage of productive T cells (bottom) between pre and post-NACT site-matched and site-unmatched samples. Paired and unpaired tests were used accordingly. TCR clonality is expressed as 1-entropy with values near 1 representing samples with one or a few predominant TCR rearrangements, while values near 0 represent more polyclonal samples D) Least absolute shrinkage and selection operator (LASSO) regression analysis using the pre-NACT matched (n=8 samples) hallmark and consensus TME cell NES as explanatory variables, and the log_2_ of the TCR clonality ratio of post/pre-NACT as response variable (see Methods). DF: Degrees of freedom. The variables selected by the LASSO regression are indicated with an asterisk. E) Spearman’s rank correlation of pre-NACT cytotoxic NES and log_2_ of the post/pre-NACT TCR clonality ratio.

Finally, we investigated whether hallmark pathways or consensus TME gene signatures calculated from the pre-treatment samples could explain the increase of TCR clonality upon neoadjuvant chemotherapy in site-matched samples (n=8, only 8 pre-treated samples with both gene expression and TCRseq data were available, see Figure 5A, Supplementary Table 5B). To perform the analysis in an unbiased manner, we employed least absolute shrinkage and selection operator (LASSO) regression analysis with the change of TCR clonality before and after NACT as a response variable (see Methods). The hallmark pathways that potentially have predictive value with positive association were the hallmarks G2M checkpoint, Myc and Wnt signaling and UV response, while apoptosis and PI3K-AKT-MTOR showed a negative association with TCR clonality increase (Figure 6D). We then performed the same analysis using the consensus TME gene signatures, where B cells, M1 macrophages and endothelial cells showed a positive association with TCR clonal expansion, while cytotoxic, CD4 and plasma cells showed a negative association. In addition, the LASSO analysis selected the consensus cytotoxic signature as relevant for explaining T cell clonal expansion upon NACT, and a correlation analysis supported this association (Figure 6E). Post-selection inference, taking into account for the uncertainty of the model selection and multiplicity, corroborated that the pre-NACT NES of the Consensus cytotoxic gene set is a promising variable to explain T cell clonal expansion upon NACT in these eight samples (*p*=0.096, see Methods). Overall, these results show that chemotherapy induces a T cell activation in HGSOC in site-matched samples but not in site-unmatched samples, further suggesting that different local immune-microenvironments play a role in the response to chemotherapy treatment. Also, pre-treatment samples with low T cell infiltration have a significant increase of TCR clonality upon chemotherapy, while tumors with higher infiltration levels do not have such a high clonal expansion, an observation that could not be addressed in unmatched tumor samples, due to the intra-patient tumor-immune heterogeneity. This highlights the potential confounding effects that intra-patient tumor-immune heterogeneity imposes on comparing tumors not only from different patients but also from different sites within the same patient.

## DISCUSSION

Despite advances in surgical approaches, chemotherapy and targeted therapies, the prognosis for patients with high grade serous ovarian cancer remains poor, with the near-inevitable development of resistance to systemic therapy. Genetic and molecular analyses of asynchronous and disseminated tumors within patients have recently started to shed light on tumor clonal dynamics and evolutionary properties of different tumor types (Johnson et al. 2014; Yates et al. 2015; McPherson et al. 2016); however, the extent of TME heterogeneity in advanced HGSOC has only begun to be revealed (Jiménez-Sánchez et al. 2017; A. W.Zhang et al. 2018). We explored the main sources of variation in the transcriptomic space among treatment-naive samples and detected that transcriptomic pathway heterogeneity is mainly explained by presence or absence of immune and stromal cells. Importantly, the degree of immune signature variation within patients was similar to the extent we observed in a case study of metastatic HGSOC, where different tumor immune microenvironments were associated with clinical outcome. In the index case, tumors with high immune related pathways regressed and presented evidence of T cell activation, while immune excluded tumors progressed (Jiménez-Sánchez et al. 2017). In the present study, all patients presented at least one tumor with low immune infiltration, suggesting that HGSOC is characterized by microenvironmental niches, which could underlie primary and acquired resistance to therapies (Wang et al. 2016; Sharma et al. 2017; Hirata et al. 2015). Through tissue image analysis, we captured immune signature differences and variation of T cell infiltration within tumors which was confirmed by immunofluorescence staining of T cells. Taken together, the transcriptional, imaged-based and immunofluorescence analyses show that TME heterogeneity is an intrinsic feature of HGSOC, which spans across patients, tumors within patients and within tumors. Furthermore, we found that intra-patient TME heterogeneity can mask the immune activation generated by treatment with cytotoxic chemotherapy. These analyses provide firm evidence that the TME affects the extent of immune activation generated upon treatment with chemotherapy.

Since the TME can constrain or foster tumor progression, targeting the TME represents a promising alternative to complement therapeutic strategies (Hansen, Coleman, and Sood 2016). However, the degree to which TME heterogeneity is driven by stochastic, cellular or molecular processes is not well understood, and little is known on the potential mechanisms behind TME spatial heterogeneity in HGSOC. Previous unbiased immune deconvolution studies have estimated relative abundances of immune cells in ovarian cancer tumors (Newman et al. 2015), and calculated survival associations (B.Li et al. 2016); however, no objective benchmarking has been performed to date, and discordant results make it difficult to judge these observations (B.Li, Liu, and Liu 2017; Newman et al. 2017; Zheng 2017b). We integrated data from different deconvolution methods and generated a consensus approach which consistently improved the predictions on our data sets and in TCGA ovarian cancer leukocyte methylation data. This showed that cytotoxic and NK cells are the major populations present in low purity tumors, while endothelial cells are the main TME cell type in high purity tumors. A previous pan-cancer analysis showed negative correlations between somatic copy number alterations (SCNA) and deconvoluted immune infiltrates, and that NK cells and CD8 T cell receptor pathway were the most differentially abundant immune factors between tumor samples with high versus low SCNA (Davoli et al. 2017). Despite these associations, the molecular mechanisms underlying this negative correlation between immune infiltrate and SCNA in HGSOC were not elucidated. Interestingly, NK cell infiltration appeared to be significantly higher in tumors with low compared to high SCNAs in HGSOC (Davoli et al. 2017), and our results also showed that NK cells were more enriched than CD8 and CD4 T cell infiltration in low purity tumors. Together, these results indicate that NK cells could be a potential TME cellular target in HGSOC, and further investigations would be required to validate this.

Transcriptional signatures can be highly informative as diagnostic/prognostic resources, as well as provide insights on mechanistic underpinnings (Burel and Peters 2018). In this study we took advantage of the availability of having multiple tumors from the same patients and performed differential expression analysis between high and low purity tumors. Pathway analysis of the differentially expressed genes showed that Wnt and Myc signaling pathways were more prevalent in purer tumors, consistent with emerging data in HGSOC and other tumors and models (Gounari et al. 2002; Damsky et al. 2011; Spranger, Bao, and Gajewski 2015; Spranger et al. 2016; Sridharan et al. 2016; Spranger et al. 2017). In addition, different studies have observed association between loss of *p53* function and decrease of NK cell infiltration in mouse models (Xue et al. 2007) and T cell infiltration human breast cancers (Iannello et al. 2013) In fact, missense or nonsense mutations in *p53* are the earliest and almost ubiquitous (96%) alterations in HGSOC (Ahmed et al. 2010; Cancer Genome Atlas Research Network 2011). Overall, these different lines of evidence help to clarify why HGSOC is intrinsically non-immunogenic: beyond the low somatic missense mutation load and the high SCNAs, the intrinsic oncogenic signaling of HGSOC seem to shape the TME and hinder immune infiltration of T cells and NK cells.

There are significant potential clinical implications from understanding the effect of chemotherapy on the TME and the molecular drivers of the heterogeneity observed, as novel combination therapies or changes in timing of treatments have the potential to improve outcomes (Patel and Minn 2018). A previous study investigated the effect of NACT on the activation of CD8, CD4 and Tregs in HGSOC, as well as systemic levels of cytokines (Böhm et al. 2016). This study found that patients who had good responses to NACT had a decrease of Tregs after treatment compared to poor responders. In general, there was also a trend towards higher cytolytic activity in tumors after NACT despite failing to detect significant changes on CD8 T cell counts (Böhm et al. 2016). Using our unbiased approach and using the consensus deconvolution method, we observed an increase of cytotoxic immunogenic activity after NACT in matched tumor samples but not in site-unmatched samples from the same patient. Similarly, employing TCRseq, we found a significant increase in T cells and TCR clonality in matched samples, but no significant difference was detected in unmatched pairs. Comparing post-NACT site-matched and post-NACT site-unmatched samples indicated that the observed change in TCR clonal expansions was driven by chemotherapy and not by the biopsy itself, although we cannot formally exclude potential immunogenic effects that the biopsy procedure may have in a neoadjuvant setting. Together, these results show a clear, confounding effect that spatial TME heterogeneity can cause.

Having unmasked the immune activation generated by NACT, we used the matched samples to investigate the factors in pre-treated samples that influence TCR clonal expansion induction upon NACT. Interestingly, cytotoxic cells showed a significant negative association with TCR clonal expansion, and the PI3K-AKT-MTOR pathway, which is part of the TCR signaling cascade, also showed a strong negative association. Conversely, Wnt and Myc signaling appeared as positively associated with TCR clonal expansion. Together, these results suggest that tumors with low levels of infiltration have a higher potential for T cell activation upon NACT than tumors with a previous immune presence. We hypothesize that this could be due to an already exhausted immune TME generated as a consequence of chronic immune and tumor interaction, while immune excluded tumors present a fresh environment where T cells can become active and expand. Importantly, our results point towards NK cells being similar or more activated after NACT, suggesting another potential therapeutic cellular target for combination therapy. Finally, this conclusion could only be drawn only when intrapatient spatial TME heterogeneity was controlled for, again highlighting the necessity to take TME heterogeneity into consideration in translational studies and in clinical applications.

Here we have presented how TME heterogeneity is a common feature of HGSOC and how that can affect the interpretation of translational studies. We have sought to uncover molecular and cellular mechanisms behind intra-patient and intra-tumor TME heterogeneity. However, there are critical limitations to consider. Disentangling the actual mechanisms using human tumor samples represents a formidable challenge since tissue samples are limited, inter-patient variability is prominent and mechanistic experimental validation is prohibitive. Given these constraints, this study is descriptive in nature and relies heavily on observations derived by independent studies using murine tumor models to propose suitable explanations. Despite this main limitation, our unbiased analysis of human tumors not only complements experimental studies, but also provides new hypotheses to further explore in a pre-clinical and clinical settings. Another major limitation is the small number of samples available compared to the large parameter space to investigate in an unbiased systematic study, which poses a challenge not only for achieving statistically meaningful results but also for the findings to be influenced by confounding variables. However, the implementation of orthogonal methods in combination with the independence of the two cohort of patients provide solid evidence of TME heterogeneity at multiple dimensions in addition to reasonable putative mechanisms behind it. Ultimately, we hope that the increasing scientific evidence would lead to better designed clinical trials where TME heterogeneity is further evaluated, data driven combination therapies or novel therapies are tested and tissue sections are systematically analyzed and stored for future integrative unbiased analyses.

This study shows that the TME of HGSOC is intrinsically heterogeneous within patients and within tumors, posing an important barrier for the successful application of therapies that target the TME, like checkpoint blockade immunotherapy. By controlling patient dependency and accounting for intra-patient TME heterogeneity, insights into potential mechanisms driving TME heterogeneity were obtained, putting forward new therapeutic strategies to be explored in future studies. Furthermore, the induced immunogenicity upon NACT treatment was only unmasked after taking into account the TME heterogeneity, which otherwise acts as a confounding variable. Despite high rates of response to initial treatment, HGSOC has a high recurrence rate and has yet to show significant response to available immunotherapeutic agents. Exploring new combination therapies and novel therapeutic targets based on a greater understanding of the TME has the potential to change the current paradigm of treatment, and hopefully improve clinical outcomes in this disease.

## METHODS

### Contact for Reagent and Resource Sharing

Further information and requests for resources and reagents should be directed to and will be fulfilled by the Lead Contact, Martin L. Miller.

### Experimental Model and Subject Details

#### Patients

All patients had stage IIIC or IV high grade serous ovarian cancer as assessed by a pathologist specialized in gynecologic malignancies. Patients signed written consent to Institutional Review Board (IRB)-approved bio-specimen protocols.

#### Treatment-naive cohort

For the treatment-naive cohort, 25 patients were consented to the study between August 2014 and March 2016. Out of these patients, 17 were excluded as they either a) withdrew from the study (n=3); b) the final pathology was not HGSOC (n=5); c) the patients had disease progression upon review of study imaging and underwent neoadjuvant chemotherapy instead of primary cytoreductive surgery (n=5); d) inadequate image-guided tissue sampling due to friable tissue (n=2); e) research imaging studies not performed due to patient cancellation (n=2). The final treatment-naive study population consisted of 8 patients with histopathologically-confirmed diagnosis of HGSOC (Supplementary Table 1A-B). Each patient underwent multi-parametric MRI (mpMRI) of the abdomen and pelvis and 18F-FDG PET/CT within 7 days immediately preceding the primary cytoreductive surgery. Volumetric regions of interest (VOI) were outlined on both axial T2-weighted MR images and PET images, covering both the primary and metastatic lesions, using ImageJ (U.S. National Institutes of Health) by a board certified radiologist with special expertise in ovarian cancer imaging. The tumor regions outlined on MRI were co-registered with those outlined on PET.

#### Neoadjuvant chemotherapy cohort

For the neoadjuvant chemotherapy cohort a previously established institutional database identified 152 patients with HGSOC who underwent neoadjuvant chemotherapy between 2008 and 2013. Of these patients, 149 went on to interval debulking surgery. 48 of these patients had adequate pre and post treatment formalin fixed paraffin embedded tissue samples available. All pretreatment specimens were obtained either through core biopsy or laparoscopic biopsy, and all post treatment specimens were obtained at the time of laparotomy for interval debulking surgery. Choice of chemotherapy was at the clinician’s discretion, but all patients in the cohort received a platinum and taxane based regimen (Supplementary Table 5A). 40 of these paired samples yielded data for analysis, 17 of these pairs were site-matched, meaning that pre-treatment and post-treatment specimens were taken from the same anatomic site, while 23 were site-unmatched. Gene expression and TCRseq data were generated for 28 and 37 pairs respectively (Supplementary Table 5B). Samples with very low TCR sequences (n=5 samples, 10 pairs) were not included in the downstream analyses as the confidence of TCR clonality is low.

### Method Details

#### Image acquisition and analysis

The quantitative diffusion parameters *D* (diffusion coefficient) and *f* (the volume fraction of the blood flowing through the microvessels) derived from the intravoxel incoherent motion (IVIM) MRI (Le Bihan et al. 1988) and dynamic contrast-enhanced (DCE) MRI (Tofts 1997) parameter Ktrans (volume transfer constant between the blood plasma and the extravascular extracellular space) were generated voxel-wise using a dedicated in-house software written in Matlab (Mathworks Inc., Natick, MA, USA). The Standardized Uptake Values (SUV) of the voxels contained within each lesion on PET were also calculated (Kinahan et al. 2009). *k*-means clustering algorithm (Carano et al. 2004) of the *D*, *f*, *K*^trans^ and SUV voxels, with the number of clusters (*k*) being fixed to *k*=3 was used to identify imaging clusters/habitats. The mean and standard deviation (mean±std. dev.) of these parameters for each cluster were calculated. To establish coherence across patients (i.e. to label each cluster with the α, β, γ greek letters, such that across patients clusters would have similar imaging features), the intra-cluster distance was calculated for each cluster. The greek letter of the clusters for each patient was assigned based on the relative value of the intra-cluster distance. Specifically, for each patient, β was assigned to the cluster which had the highest intra-cluster distance for that patient; γ was assigned to the cluster with intermediate intra-cluster distance, and α to the cluster with the lowest intra-cluster distance.

#### Custom made 3D Tumor Moulds

For each patient, custom made 3 dimensional (3D) moulds [REF:Weigelt B, Vargas AH, Selenica P, Geyer FC, Mazaheri Y, Blecua P, Conlon N, Hoang LN, Jungbluth AA, Snyder A, Ng CKY, Papanastasiou AD, Sosa RE, Soslow RA, Chi DS, Gardner GJ, Shen R, Reis-Filho JS, Sala E. Radiogenomics analysis of intra-tumor heterogeneity in high-grade serous ovarian cancer. BJC (under review).] were printed based on manual segmentation of the ovarian mass and metastatic implants on axial T2-weighted MR images. The lesions were outlined on every axial slice and automatically converted into 3D models using open source software (MIPAV, National Institutes for Health, Center for Information Technology). The final 3D models of each lesion were imported into OpenSCAD (OpenSCAD, The OpenSCAD Developers), a 3D CAD modeling software, which was used to create an internal cavity that exactly shaped each lesion according to the MRI shape and contour. The custom-made 3D tumor moulds were printed using a 3D printer (MakerBot Replicator 2, MakerBot, Brooklyn, NY). The slits for slicing each lesion were placed and labelled into the molds at 10mm intervals corresponding to the slice thickness and locations of the axial T2W-weighted MR images. The mould was also labeled with left, right, anterior, posterior, superior and inferior markers to allow for proper orientation when collecting samples in the operating room.

#### Cluster guided specimen sampling

All 3D moulds containing the specimens were taken to the histopathology department where each lesion was sampled by a pathology fellow. Each tumor was sectioned through the mould and samples were taken according to the imaging habitats/clusters defined above. Half of the sample was sent for histopathology and the other one for immunogenomic analysis.

#### Immunofluorescent Staining

The immunofluorescent staining was performed at Molecular Cytology Core Facility of Memorial Sloan Kettering Cancer Center using Discovery XT processor (Ventana Medical Systems). The tissue sections were deparaffinized with EZPrep buffer (Ventana Medical Systems), antigen retrieval was performed with CC1 buffer (Ventana Medical Systems). Sections were blocked for 30 minutes with Background Buster solution (Innovex), followed by avidin-biotin blocking for 8 minutes (Ventana Medical Systems).

Multiplex immunofluorescence stainings were performed as previously described (Yarilin et al. 2015). Slides were incubated with anti-FoxP3 (Abcam, cat#ab20034, 5 ug/ml) for 4 hours, followed by 60 minutes incubation with biotinylated horse anti-mouse IgG (Vector Labs, cat# MKB-22258) at 1:200 dilution. The detection was performed with Streptavidin-HRP D (part of DABMap kit, Ventana Medical Systems), followed by incubation with Tyramide Alexa Fluor 488 (Invitrogen, cat# T20922) prepared according to manufacturer instruction with predetermined dilutions. Next, sections were incubated with anti-CD4 (Ventana, cat#790-4423, 0.5ug/ml) for 5 hours, followed by 60 minutes incubation with biotinylated goat anti-rabbit IgG (Vector, cat # PK6101) at 1:200 dilution. The detection was performed with Streptavidin-HRP D (part of DABMap kit, Ventana Medical Systems), followed by incubation with Tyramide Alexa 568 (Invitrogen, cat# T20914) prepared according to manufacturer instruction with predetermined dilutions. Finally, sections were incubated with anti-CD8 (Ventana, cat#790-4460, 0.07ug/ml) for 5 hours, followed by 60 minutes incubation with biotinylated goat anti-rabbit IgG (Vector, cat # PK6101) at 1:200 dilution. The detection was performed with Streptavidin-HRP D (part of DABMap kit, Ventana Medical Systems), followed by incubation with Tyramide Alexa 647 (Invitrogen, cat# T20936) prepared according to manufacturer instruction with predetermined dilutions. After staining slides were counterstained with DAPI (Sigma Aldrich, cat# D9542, 5 ug/ml) for 10 min and coverslipped with Mowiol.

Stained slides were digitized using Pannoramic Flash 250 (3DHistech, Hungary) using 20x/0.8NA objective. Regions of interest were drawn on the scanned images using Pannoramic Viewer (3DHistech, Hungary) and exported into tiff images. ImageJ/FIJI was used to segment DAPI-stained nuclei and count the cells with positive signal.

#### Nucleic Acid Isolation and Quantification

DNA and RNA were extracted from tumor areas delineated as tumor nests on H&E slides reviewed by a pathologist specialized in gynecologic malignancies using the DNeasy^®^ and RNeasy^®^ (Qiagen) assays, respectively. RNA expression was assessed using the human Affymetrix Clariom D Pico assay (Thermo Fisher Scientific).

#### T-Cell Receptor Sequencing

High-throughput in-situ sequencing of the T cell receptors present in the samples and blood of the patient was performed using the immunoSEQ assay platform (Adaptive Biotechnologies).

### Quantification and Statistical Analysis

#### Gene expression analysis

RNA expression was assessed using the human Affymetrix Clariom D Pico assay. Arrays were analyzed using the SST-RMA algorithm in the Affymetrix Expression Console Software. Expression was determined by using the Affymetrix Transcriptome Analysis Console. Locally weighted scatterplot smoothing (LOESS) normalization across samples was implemented (Gautier et al. 2004) using:

~~~
# R 3.5.0
library(affy) # version 1.58.0
data_norm<-normalize.loess(data, family.loess=‘gaussian’)
~~~

#### Single-sample gene set enrichment analysis

Single-sample gene set enrichment analysis (Barbie et al. 2009), a modification of standard GSEA (Subramanian et al. 2005), was performed on RNA measurements for each sample using the GSVA package version 1.28.0 (Hänzelmann, Castelo, and Guinney 2013) in R version 3.5.0 with parameters: `method = ‘ssgsea’`, and `tau = 0.25.` Normalized enrichment scores were generated for the hallmark gene sets (Arthur Liberzon et al. 2015), immune and stromal signatures (Yoshihara et al. 2013), TME cell gene sets obtained from previous publications (Bindea et al. 2013; Davoli et al. 2017), as well as the consensus TME gene sets (Figure S3A). Hallmark gene sets were obtained from MSigDB database version 6.1 (A.Liberzon et al. 2011).

#### Tumor purity and immune cell gene-expression score

Tumor purity and total immune component in the tumor samples were analyzed using the ESTIMATE algorithm method version 1.0.13 (Yoshihara et al. 2013) on the gene expression data using the option: `platform = ‘affymetrix’` for the cohort samples and `platform = ‘illumina’` for TCGA OV samples, in R version 3.5.0.

#### Dimensionality reduction

The t-distributed Stochastic Neighbor Embedding (t-SNE) and principal component analysis dimensionality reduction methods were implemented in `python version 3.6.5` using the `sklearn.manifold.TSNE` and `sklearn.decomposition.PCA` functions from the `sklearn version 0.19.1` package (Pedregosa et al. 2011). PCA was computed after sample wise standardization. Functions were used as follows:

~~~
X_tsne=tsne.TSNE(learning_rate=100,n_iter=5000,perplexity=5).fit_trans form(loess_normalized_expression.values)
X_pca=PCA(n_components=7).fit_transform(nes_standarized)
~~~

#### Analysis of T cell infiltration between cases, sites, and habitats

A linear mixed effects model analysis was performed to evaluate if there were significant differences in T cell infiltration subsets between patients, sites within patients, and habitats within tumors, and to assess whether the differences were other than random. Due to data nested dependency we employed a linear mixed effects model under the lme4 R package (Bates et al. 2015). Cases were considered a random factor, while sites and habitats were considered fixed factors as follows:

~~~
# R 3.5.0
library(lme4) # version 1.1-17
glmer.fit1a <- glmer(cbind(cd8_counts, total_cells - cd8_counts) ~
site*habitats+(1|case), family = binomial, data = data)
~~~

#### TME cell deconvolution methods

Cell deconvolution methods were used to estimate levels of non-cancerous cells in the TME. The methods employed were CIBERSORT (Newman et al. 2015), MCP-counter (Becht et al. 2016), TIMER (B.Li et al. 2016), xCELL (Aran, Hu, and Butte 2017), as well as gene sets collected from two previous publications (Bindea et al. 2013; Davoli et al. 2017).

CIBERSORT analysis was performed using the CIBERSORT R script as follows:

~~~
# R 3.5.0
source(’CIBERSORT.R’) # version 1.04
cibersort <- CIBERSORT(’LM22.txt’,’expression_data.txt’, perm=1000,
QN=TRUE, absolute=FALSE)
~~~

MCP-counter analysis was performed as follows:

~~~
# R 3.5.0
library(MCPcounter) # version 1.1.0
exp_data = read.table(expression_data.txt, header=T, sep=’\t’,
row.names=’Hugo_Symbol’)
mcp = MCPcounter.estimate(exp_data,
featuresType=c(“affy133P2_probesets”,“HUGO_symbols”,“ENTREZ_ID”)[2],
probesets=read.table(curl(“http://raw.githubusercontent.com/ebecht/MCPcounter/master/Signatures/probesets.txt”),sep=“\t”,stringsAsFactors=FALSE,colClasses =“character”),
genes=read.table(curl(“http://raw.githubusercontent.com/ebecht/MCPcounter/master/Signatures/genes.txt”),sep=“\t”,stringsAsFactors=FALSE,header=TRUE,colCla sses=“character”,check.names=FALSE))
~~~

The TIMER web server (https://cistrome.shinyapps.io/timer/) was used for deconvolution of TME cells (T. Li et al. 2017).

The xCELL web server version 1.1 (http://xcell.ucsf.edu/) was used for deconvolution of TME cells.

For the Bindea *et al*. and Davoli *et al*. gene sets, standard ssGSEA analysis was performed as previously described.

#### Consensus TME cell gene sets

To generate the consensus TME gene sets we identified cell types that were deconvoluted by at least 2 different methods, leading to 18 different cell types. We then intersected the gene sets that the different methods considered for the deconvolution of such cell types. To intersect genes used in CIBERSORT, we first filtered out genes whose weight was below 1.96 standard deviations of the mean for each of CIBERSORT cell types. In addition we combined activated states into the corresponding cell type. The only activated stated included was cytotoxic cells, which would include CD8 and/or NK cells in their activated stated. The intersected genes were used to represent each cell type, and genes with a higher Pearson’s correlation coefficient than −0.2 and a *p*-value ≤ 0.05 with tumor purity as defined by TIMER were filtered out from the gene sets (B.Li et al. 2016). Finally, ssGSEA was employed to calculate NES for each cell type as described above (Figure S3A).

#### TME cell deconvolution benchmarks

##### T cell subsets immunofluorescent staining benchmark

We correlated the CD8, CD4, Tregs infiltration counts with the deconvolution scores generated by ESTIMATE, CIBERSORT, MCP-counter, Bindea *et al*., Davoli *et al*., TIMER, xCELL, and the consensus TME scores. For the immune score comparison, all the genes used for the deconvolution for each method were aggregated together into one single gene set per method except for CIBERSORT. CIBERSORT deconvolution-log_10_(*p*-values) were used as a metric for immune score comparison. CD8, CD4, and Treg counts from IF data were summed and used for the comparison. Because the methods have different scoring systems and ranges we standardized (z-score) the scores to be able to compare the results across methods together. For each tumor, multiple IF-stained sections were quantified for TILs, and we correlated all the regions quantified with the deconvolution scores of each tumor, explaining the vertical patterns observed in figure 3A. Spearman’s rank correlation was performed for each comparison and FDR *p*-value correction was applied.

##### TCGA OV leukocyte methylation benchmark

As an independent benchmark we used leukocyte methylation scores of TCGA ovarian cancer samples (Cancer Genome Atlas Research Network 2011). TCGA ovarian cancer RNAseq data was retrieved from the cBioPortal (Cerami et al. 2012; Gao et al. 2013) web server version 1.6.2 (http://www.cbioportal.org/).

First, deconvolution of cell types was performed using the different methods listed above. Spearman’s rank correlations were calculated between CD8 T cell scores and the leukocyte methylation score of TCGA ovarian cancer samples, and FDR *p*-value correction was applied.

We further fitted multiple linear regression models to each method deconvoluted cell types (Figure S3B). We compared the proportion of leukocyte methylation score variance that is explained by the unsupervised selected immune cells (adjusted R-squared), as well as the relative quality of the models by considering goodness of fit and model simplicity after BoxCox transformation of the leukocyte methylation scores to meet the normality of residuals assumption. As a sensitivity analysis we log transformed the leukocyte methylation scores before performing the linear regression models. In both analyses (BoxCox and log-transform), stepwise Akaike information criterion variable selection was performed once normality and heteroscedasticity assumptions were checked. Akaike information criterion (AIC) and Bayesian information criterion (BIC) were employed independently to compare the fitted models for each method. Both AIC and BIC quantify information loss and penalize the number of variables. Thus, the trade-off between goodness of fit and model simplicity across methods was evaluated, allowing us to quantify and minimize information loss. The R version and packages used for this analysis were R version 3.5.0, gamlss_5.0-8 (Stasinopoulos, Mikis Stasinopoulos, and Rigby 2007), leaps_3.0, car_3.0-0 (Fox and Weisberg 2011), and MASS_7.3-50 (Venables and Ripley 2002).

#### Differential expression analysis

Tumor samples from the treatment-naive cohort were divided into high and low purity classes taking as a cutoff the median of tumor purity calculated for the tumor samples using ESTIMATE (Yoshihara et al. 2013). Then a differential expression analysis was performed using the R packages limma_3.36.1 (Ritchie et al. 2015) and Biobase_2.40.0 (Huber et al. 2015). Patient dependency was taken into account as follows:

~~~
# R 3.5.0
library(limma) # version limma_3.36.1
library(Biobase) # version Biobase_2.40.0
data<- read.table(’expression_data.txt’,sep=’\t’,header=T,row.names=’Hugo_Symbol’)
eset<-new("ExpressionSet", exprs=as.matrix(data))
estimate<-
read.table(’estimate_purity_scores’,sep=’\t’,header=T,row.names=’NAME’)
med_purity<-median(estimate$TumorPurity)
purity<-as.data.frame(ifelse(estimate$TumorPurity > med_purity,
‘high_purity’, ‘low_purity’))
row.names(purity)<-row.names(estimate)
colnames(purity)[1]<-’purity’
patient_data<-
read.table(’patient_data.txt’,sep=’\t’,header=T,row.names=’NAME’)
clinical<-merge(patient_data,purity,by=’row.names’)
row.names(clinical)<-row.names(purity)
clinical$Row.names<-NULL
colnames(clinical)[6]<-’purity’
# merge factors
clinical<-factor(clinical$purity)
# Make design matrix
design <- model.matrix(~0+clinical)
colnames(design) <- levels(clinical)
# estimate correlation between measurements on same subjects
corfit <- duplicateCorrelation(eset,design,block=patient_data$case)
# inter-subject correlation is input into the linear model fit
fit <-
lmFit(eset,design,block=patient_data$case,correlation=corfit$consensus)
cm <- makeContrasts(HighPurityvsLowPurity = high_purity-low_purity,
levels=design)
fit2 <- contrasts.fit(fit, cm)
fit2 <- eBayes(fit2)
results <- decideTests(fit2, adjust.method="fdr")
volcanoplot(fit2)
~~~

#### Gene ontology analysis

Gene ontology analysis of significantly up or down-regulated genes was performed using the Gene Ontology Consortium (Ashburner et al. 2000; The Gene Ontology Consortium 2017) web server (http://www.geneontology.org/). *P*-value FDR corrections were calculated for this analysis.

#### ssGSEA of differential expression analysis

Further, *p*-values for each gene were retrieved and multiplied by −1 if the the log_2_ change was negative. The list of genes with their associated *p*-value was used to calculate hallmark and consensus TME normalized enrichment scores (NES) through ssGSEA. Hallmark gene sets’ NES were normalized by taking the exponential function. Consensus TME gene sets’ NES approached normality by taking the natural logarithm. Modified z-score was employed to detect outliers in the hallmarks and consensus TME NES independently, as the modified z-score uses the median and the median absolute deviation (MAD) to robustly measure central tendency and dispersion in small data sets (Iglewicz and Hoaglin 1993).

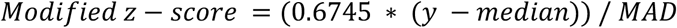

#### Paired gene set comparisons

##### Volcano plots

For each of the 52 hallmark and 18 consensus TME gene sets paired comparisons before and after NACT were performed. Equality of variance (Bartlett’s test) and normality (Shapiro test, Kolmogorov-Smirnov test, and D-Agostino-Pearson’s test) assumptions were checked to select the corresponding paired test (Paired T-test, Welch’s T-test, or Wilcoxon signed-rank test). The analysis was performed under python 3.6.5 and scipy 1.1.0 (http://www.scipy.org/) ecosystem (Oliphant 2007).

##### Hotelling’s T^2^ distribution test

Multivariate Hotelling’s T^2^ test was performed to compare difference of CD8, NK, and cytotoxic consensus TME gene sets NES between pre- and post-NACT tumors as follows:

~~~
# R 3.5.0
library(Hotelling) # version Hotelling_1.0-4
data<-read.table(’consensusTME_NES.txt’,sep=’\t’,header=T,row.names=’NAME’)
matched<-data[which(data$related==’matched’),]
matched<-subset(matched, select = -c(case_number,Site,related))
unmatched<-data[which(data$related==unmatched),]
unmatched<-subset(paired, select = -c(case_number,Site,related))
fit_matched = hotelling.test(nk*cd8*cytotoxic~NACT, data=matched, perm=T)
fit_unmatched = hotelling.test(nk*cd8*cytotoxic~NACT, data=unmatched, perm=T)
plot(fit_matched)
plot(fit_unmatched)
fit_matched$pval
fit_unmatched$pval
~~~

#### TCR sequencing analysis

Analysis of the sequences was performed on the immunoSEQ ANALYZER 3.0 (Adaptive biotechnologies). T cell counts and TCR clonality were retrieved for statistical comparisons. T cell counts are derived from quantitative immunoSequencing of the TCRB loci, in which the internal controls allow precise quantitation of sequence counts based on reads. Nucleated cell counts are determined by sequencing housekeeping genes. The fraction of T cells is determined by dividing the T cell count by the nucleated cell counts. Values for TCR productive clonality range from 0 to 1. Values near 1 represent samples with one or a few predominant rearrangements (monoclonal or oligoclonal samples) dominating the observed repertoire. TCR productive clonality values near 0 represent more polyclonal samples. TCR productive clonality is calculated by normalizing productive entropy using the total number of unique productive rearrangements and subtracting the result from 1.

#### LASSO regression and post-selection inference

Least absolute shrinkage and selection operator (LASSO) regression analysis was performed using the glmnet R package (Friedman, Hastie, and Tibshirani 2010). Hallmark and consensus TME cell type NES of pre-NACT samples were used independently as explanatory variables, and the log_2_ of the ratio post/pre NACT TCR clonality as response variable.

~~~
# R 3.5.0
library(glmnet) # version glmnet_2.0-16
dataH<-
read.table(’Hallmarks_TCRdiff.txt’,sep=’\t’,header=T,row.names=’ID’)
dataC<-
read.table(’ConsensusTME_TCRdiff.txt’,sep=’\t’,header=T,row.names=’ID’)
h<-as.matrix(subset(dataH, select = -c(clon_dif)))
c<-as.matrix(subset(dataC, select = -c(clon_dif)))
y<-as.matrix(dataH$clon_dif)
hallmark_fit<-glmnet(h,y,family=’gaussian’,alpha=1,standardize=T)
tmecells_fit<-glmnet(c,y,family=’gaussian’,alpha=1,standardize=T)
# VARIABLE SELECTION
hallmark_cvfit=cv.glmnet(h,y,family=“gaussian”,type.measure=“mse”,alpha=1,
nfold=10)
tmecells_cvfit=cv.glmnet(c,y,family=“gaussian”,type.measure=“mse”,alpha=1,
nfold=10)
coef(hallmark_cvfit, s = “lambda.min”)
coef(tmecells_cvfit, s = “lambda.min”)
# POST-SELECTION INFERENCE
library("selectiveInference") # version selectiveInference_1.2.4
postselinf_h = fixedLassoInf(h,y,beta_hat[-1],lambda,family=“gaussian”)
postselinf_c = fixedLassoInf(c,y,beta_hat[-1],lambda,family=“gaussian”)
~~~

## Data and Software Availability

Requests for additional data and custom code should be directed to the corresponding authors.

### Immunofluorescence staining images

The immunofluorescence images discussed in this study will be provided upon request to the Lead Contact in the Data and Software Availability section.

### Microarray data

Microarray data is currently under submission to the GEO database.

### TCR sequencing data

The TCR sequencing data discussed in this study will be provided upon request to the Lead Contact in the Data and Software Availability section.

## ACKNOWLEDGEMENTS

We acknowledge Tony Wu for his insightful comments on the manuscript. A.S. was supported by grants from the Marsha Rivkin Organization, Society of Memorial Sloan Kettering, Translational and Integrative Medicine Research Fund (MSKCC), Kaleidoscope of Hope, and a MSKCC Cancer Center Support Grant of the National Institutes of Health/National Cancer Institute (P30 CA008748). M.L.M. was supported by Cancer Research UK core grant (C14303/A17197). A.J.S. was supported by a doctoral fellowship from the Cancer Research UK Cambridge Institute and the Mexican National Council of Science and Technology (CONACyT).

## CONTRIBUTIONS

*CRediT standard taxonomy*

http://dictionary.casrai.org/Contributor_Roles

**Conceptualization** [Ideas; formulation or evolution of overarching research goals and aims]

AS, AJS, MLM, ES

**Data curation** [Management activities to annotate (produce metadata), scrub data and maintain research data (including software code, where necessary for interpreting the data itself) for initial use and later re-use]

AJS, KL, PC

**Formal analysis** [Application of statistical, mathematical, computational, or other formal techniques to analyse or synthesize study data]

AJS

**Funding acquisition** [Acquisition of the financial support for the project leading to this publication]

AV, ES, AS, MLM

**Investigation** [Conducting a research and investigation process, specifically performing the experiments, or data/evidence collection]

PC, KL, TW, YM, IN, BW, DC, ES

**Methodology** [Development or design of methodology; creation of models]

ES

**Project administration** [Management and coordination responsibility for the research activity planning and execution]

AS, MLM, ES

**Resources** [Provision of study materials, reagents, materials, patients, laboratory samples, animals, instrumentation, computing resources, or other analysis tools]

AS, MLM, ES

**Software** [Programming, software development; designing computer programs, implementation of the computer code and supporting algorithms; testing of existing code components]

AJS, ES (the radiologic background)

**Supervision** [Oversight and leadership responsibility for the research activity planning and execution, including mentorship external to the core team]

AS, MLM, ES

**Validation** [Verification, whether as a part of the activity or separate, of the overall replication/reproducibility of results/experiments and other research outputs]

AJS, DLC

**Visualization** [Preparation, creation and/or presentation of the published work, specifically visualization/data presentation]

AJS

**Writing original draft** [Preparation, creation and/or presentation of the published work, specifically writing the initial draft]

AJS, AS, MLM

**Writing review & editing** [Preparation, creation and/or presentation of the published work by those from the original research group, specifically critical review, commentary or revision]

AJS, MLM, AS, KL, ES, JDB

